# Stytra: an open-source, integrated system for stimulation, tracking and closed-loop behavioral experiments

**DOI:** 10.1101/492553

**Authors:** Vilim Štih, Luigi Petrucco, Andreas M. Kist, Ruben Portugues

## Abstract

We present Stytra, a flexible, open-source software package, written in Python, that we designed to cover all the general requirements involved in larval zebrafish behavioral experiments. It allows timed stimulus presentation, interfacing with external devices and simultaneous real-time tracking of position, tail and eye motion in both freely-swimming and head-restrained preparations. It logs in standardized formats all recorded quantities, metadata, and code version to allow full provenance tracking, from data acquisition through analysis to publication. The package is modular and expandable for different experimental protocols and setups. Current releases can be found at https://github.com/portugueslab/stytra. We also provide complete documentation with examples for extending the package to new stimuli and hardware, as well as a schema and parts list for behavioral setups. We showcase Stytra by reproducing two previously behavioral protocols, one with head-restrained and the other with freely-swimming larvae, as well as an experiment in which the software is used in the context of a calcium imaging experiment, where it can trigger and communicate with other acquisition devices. Our aims are to help laboratories with little or no experience in the field to start implementing behavioral experiments and to provide a platform for sharing stimulus protocols to enable easy reproduction of experiments and straightforward validation. In addition, Stytra can easily serve as a base platform to design behavioral experiments in other model organisms.

## Introduction

The central goal of systems neuroscience is to explain the neural underpinnings of behavior. To investigate the link between sensory input, brain activity and animal behavior, relevant behavioral variables have to be recorded and quantified. Moreover, the same experimental paradigm has to be reproduced in different experimental setups in order to be combined with different recording or stimulation techniques, and it needs to be reproducible across different laboratories. However, the setups generally rely on heterogeneous hardware and custom-made software tailored to the specific requirements of one experimental apparatus. Often, the code used is based on expensive software packages (such as LabView or Matlab), with open-source options for hardware control generally limited to one particular type or brand of devices. As a consequence, the same experimental protocol has to be implemented many times, thus wasting time and increasing potential sources of error. This makes sharing the code for replicating a scientific finding under the same experimental conditions very difficult.

To address these problems, we developed Stytra, a package that encompasses all the requirements of hardware control, stimulation and behavioral tracking that we encounter in our everyday experimental work. Our system, written in pure Python, provides a framework to assemble an experiment combining different input and output hardware and algorithms for online behavioral tracking and closed loop stimulation. It is highly modular and extendable to support new hardware devices or tracking algorithms. It facilitates reuse of different components of the package, encourages building upon existing work and enforces consistent data management. The definition of experimental protocols in high-level Python scripts makes it very suitable for version control and code sharing across laboratories, facilitating reproducibility and collaboration between scientists. Finally, it runs on all common desktop operating systems (Windows, Linux and MacOS), therefore incurring no additional costs on the software side. It combines the main advantages of asynchronous dataflow processing packages like Bonsai [1] with the easy prototyping of stimuli and experimental paradigms of libraries such as Psychophysics Toolbox [2].

Stytra was developed primarily in the context of a laboratory working with larval zebrafish, and it fulfills the common requirements of behavioral paradigms used with this animal [3]. Specifically, a large part of the package provides tools for quick assembly of common visual stimuli and another provides functions for tracking eyes and tail of freely-swimming or head-restrained fish. The tracking functions consist of efficient re-implementations of published algorithms, as well as newly-developed methods. We make our hardware solutions open-source as well [4]: hardware designs provided along with the documentation describe in detail apparatuses for performing common behavioral experiments. The library could be extended to offer support for behavioral paradigms in use with different species, potentially offering a unified platform to build and share experiments in neuroscientific and behavioral research.

## Design and Implementation

### Overview and library structure

We developed Stytra using the Python programming language. We endeavored to follow best practices in software engineering: an object-oriented design, separation of user interface and data processing code, and modularity. In Stytra, new experiments can be designed using a very simple Python syntax, allowing even beginners to develop their own stimulation paradigms. Once defined, a script is controlled through a graphical user interface which can be used with no knowledge of Python. At the core of the Stytra package lies the `Experiment` object, where all the components that may be used in an experiment - from the monitor, to a camera, to the image analysis functions - are added in a modular way.

This organization enables composing different experimental paradigms with full code reuse. Improvement of different modules (*e.g.* the user interface, plotting or tracking) is therefore reflected in all experimental setups, and support for a new piece of hardware or tracking function can be added with minimal effort and interference with other parts of the project. Online image processing is organized along a sequence of steps: first, images are acquired from the camera, then the image is filtered and tracked and the tracking results are saved. Acquisition, filtering and data saving occur in separate processes. This approach improves the reliability and the performance of online behavioral tracking, and exploits the advantages of multi-core processors. After processing, streaming numerical data (such as tracking results and dynamic parameters of stimuli) is passed into data accumulators in the main thread, and a user-selected subset can be plotted in real time and saved in one of the several supported formats (Fig 1). Moreover, for every experimental session all changeable properties impacting the execution of the experiment are recorded and saved. Finally, as the software package is version-controlled, the version of the software used is saved as well, ensuring the complete reproducibility of the experiment.

**Fig 1.**
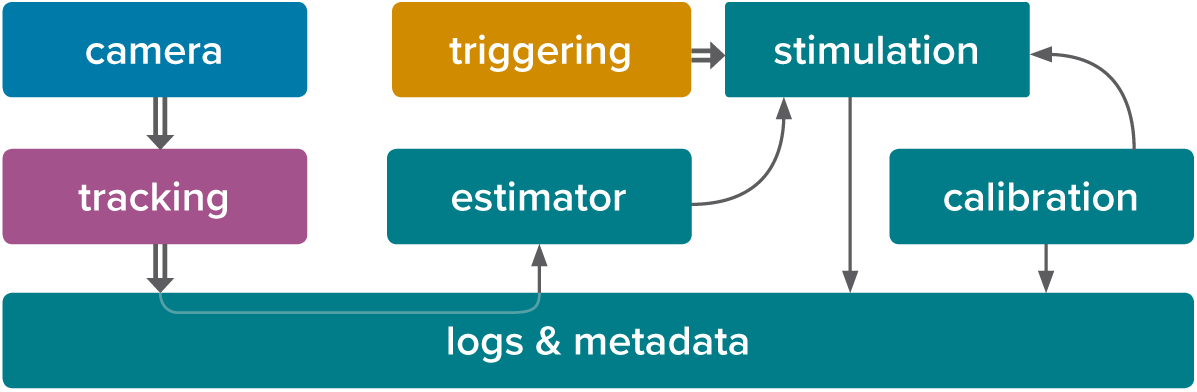
The software architecture of the system. Communication between different parts of a Stytra experiment. Each color represents a separate process in which the module(s) are running. Data flow between modules is depicted by arrows, and communication between processes is depicted as double arrows. The user interface, the stimulus update and related functions such as the screen calibration and data saving are performed in the main process, colored in green. The stimulation can be triggered by a triggering process (in yellow) that listens for an external triggering signal. Frames can be acquired from a camera process (in blue), analyzed by a tracking function (in purple), and the result can be streamed to the main process for data saving and used in closed loop-experiments via the estimator.

### Building and running an experiment in Stytra

The `Experiment` object binds all the different components required in an experiment. The most basic `Experiment` object performs the presentation of a succession of stimuli, saving the experiment metadata and the stimulation log. Subclasses like `TrackingExperiment` add features such as acquiring frames from a camera and performing online image analysis of the posture or position of the fish. Each `Experiment` is associated with a user interface to run the stimulation protocol, insert metadata, control parameters, and calibrate the stimulus display. The `Experiment` object has a protocol associated that contains the definition of the stimulus sequence and is customized by the user in a script. The appropriate `Experiment` subclass is automatically instantiated in Stytra based on the configuration defined in the protocol. For an example of how to create and run an experiment in Stytra, see the Usage examples box and the more detailed examples gallery in the documentation.

### Stimulus design

Experimental protocols in Stytra are defined as sequences of timed stimuli presented to the animal through a projector or external actuators. Each `Stimulus` object controls a specific event over time. A sequence of stimuli, defined as a Python list of `Stimulus` objects, defines a protocol (see Usage examples box). This structure makes the creation of new experimental protocols straightforward, requiring very little knowledge of the general structure of the library and only basic Python syntax. A dedicated class coordinates the timed execution of the protocol relying on a `QTimer` from the PyQt5 library, ensuring a temporal resolution in the order of 5-10 ms (around the response time of a normal monitor). Milli- or microsecond resolution, which might be required for optogenetic experiments, for example, is currently not supported. Each `Stimulus` has methods which are called at starting time or at every subsequent time step while it is set. In this way one can generate dynamically changing stimuli, or trigger external devices. New `Stimulus` types can be easily added to the library just by subclassing `Stimulus` and re-defining the `Stimulus.start()` and `Stimulus.update()` methods.

A large number of stimuli are included in the package. In particular, a library of visual stimuli has been implemented as `VisualStimulus` objects using the `QPainter` API, a part of the Qt GUI library, enabling efficient drawing with OpenGL. Relying on a high-level drawing library makes the code very readable and maintainable. The library already includes common stimuli used in visual neuroscience, such as moving bars, dots, whole-field translation or rotations of patterns on a screen, and additional features like movie playback and presentation of images from a file. The classes describing visual stimuli can be combined, and new stimuli with these patterns moved or masked can be quickly defined by combining the appropriate Stimulus types. Finally, new `VisualStimulus` objects can be defined by defining their `VisualStimulus.paint()` method. Visual stimuli are usually displayed on a second screen, therefore Stytra provides a convenient interface for positioning and calibrating the stimulation window (visible in Fig 2 on the right-hand side). Importantly, all stimulus parameters are specified in physical units and are therefore independent of the display hardware. Finally, the timed execution of code by a `Stimulus` object can be used to control hardware via I/O boards or interacting via serial communication with a MicroPython PyBoard or other microcontrollers such as the Arduino. Examples are provided together with the code (https://github.com/portugueslab/stytra).

**Fig 2.**
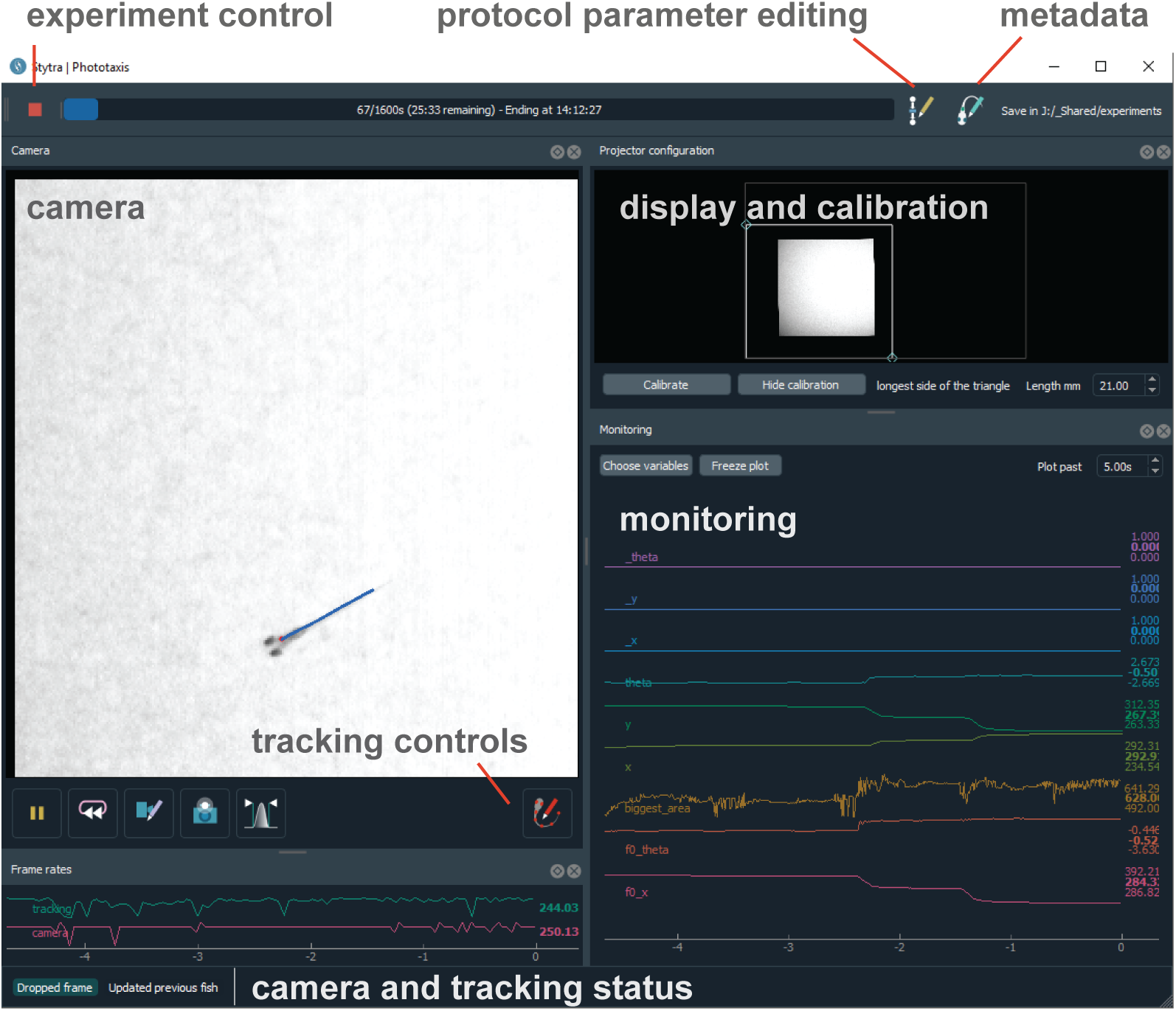
Screen capture of the software in use. The various behavioral paradigms supported by Stytra provide the user with a consistent interface to control experiments. The toolbar on top controls aspects of running the experiment, a camera panel shows the tracking results supperimposed on the camera image, a calibration panel enables quick positioning and calibration of the stimulus display and a monitoring panel plots a user-selected subset of experimental variables.

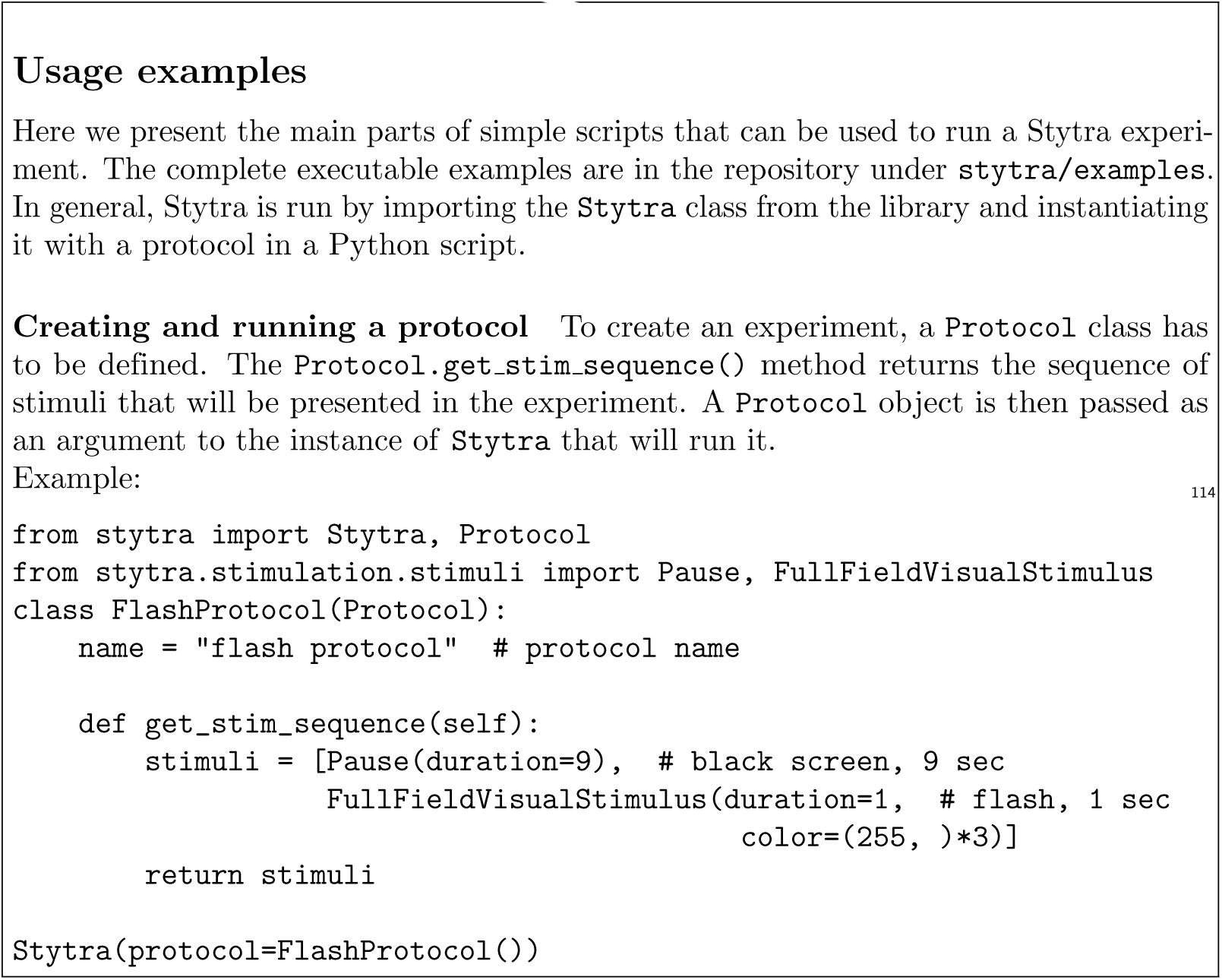

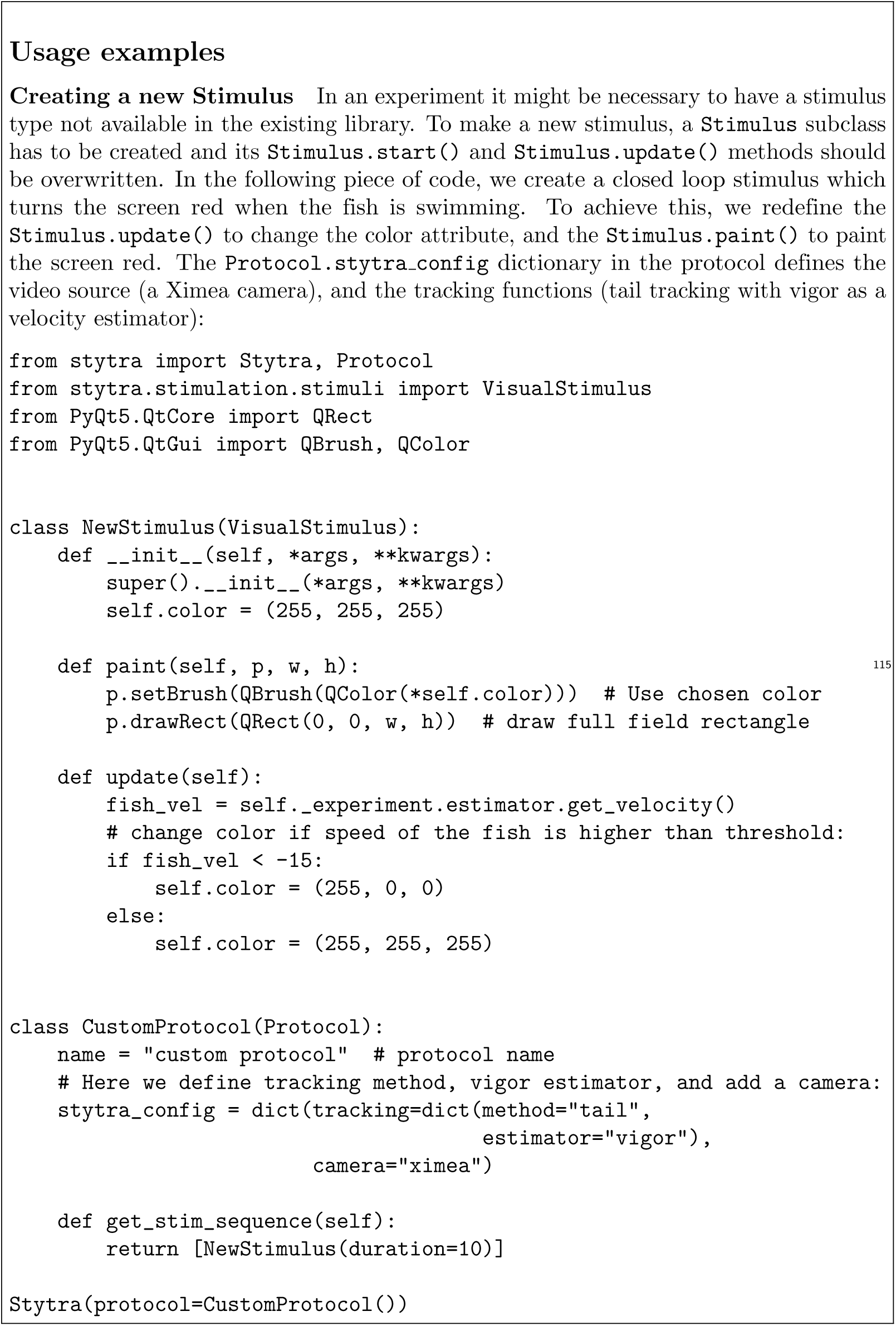

### Image acquisition and tracking

#### Image acquisition

A key feature of Stytra is the extraction of relevant behavioral features in real time from video inputs. The `Camera` object provides an interface for grabbing frames and setting parameters for a range of different camera types. Currently supported models include those by XIMEA, AVT, PointGray/FLIR and Mikrotron but others can be added as long as a Python or C API exists. Moreover, previously-recorded videos can also be processed, allowing for offline tracking. Frames are acquired from the original source in a process separated from the user interface and stimulus display. This ensures that the acquisition and tracking frame rate are independent of the stimulus display.

#### Image processing pipeline

Fish coordinates and posture are tracked by several modules through multiple processes (represented in Fig 1). A dedicated process receives frames from the image acquisition process, applies a tracking function (as well as pre-processing functions for smoothing or background subtraction), and streams its results to the main process to save and visualize the results online. This modular structure allows easy expansion of the library: new functions for pre-filtering or tracking can be incorporated into the pipeline with minimal effort. Functions to track tail and eye position in embedded fish, as well as fish position and orientation in an open arena, are already available in the tracking module. Existing functions are written using the OpenCV [5] computer vision library. In addition, custom functions are compiled with the Numba library to increase their performance.

#### Posture tracking in embedded fish

##### Tail tracking

Zebrafish larvae swim by producing alternating contractions of the tail muscles, and different types of swim bouts, from startle responses to forward swimming, require different sequences of contractions [3]. The tail of the larvae can be easily skeletonized and described as a curve discretized into 7-10 segments [6] (Fig 3A). The tail tracking functions work by finding the angle of a tail segment given the position and the orientation of the previous one. The starting position of the tail, as well as a rough tail orientation and length need to be specified beforehand using start and end points, movable over the camera image displayed in the user interface (as can be seen in Fig 2A).

**Fig 3.**
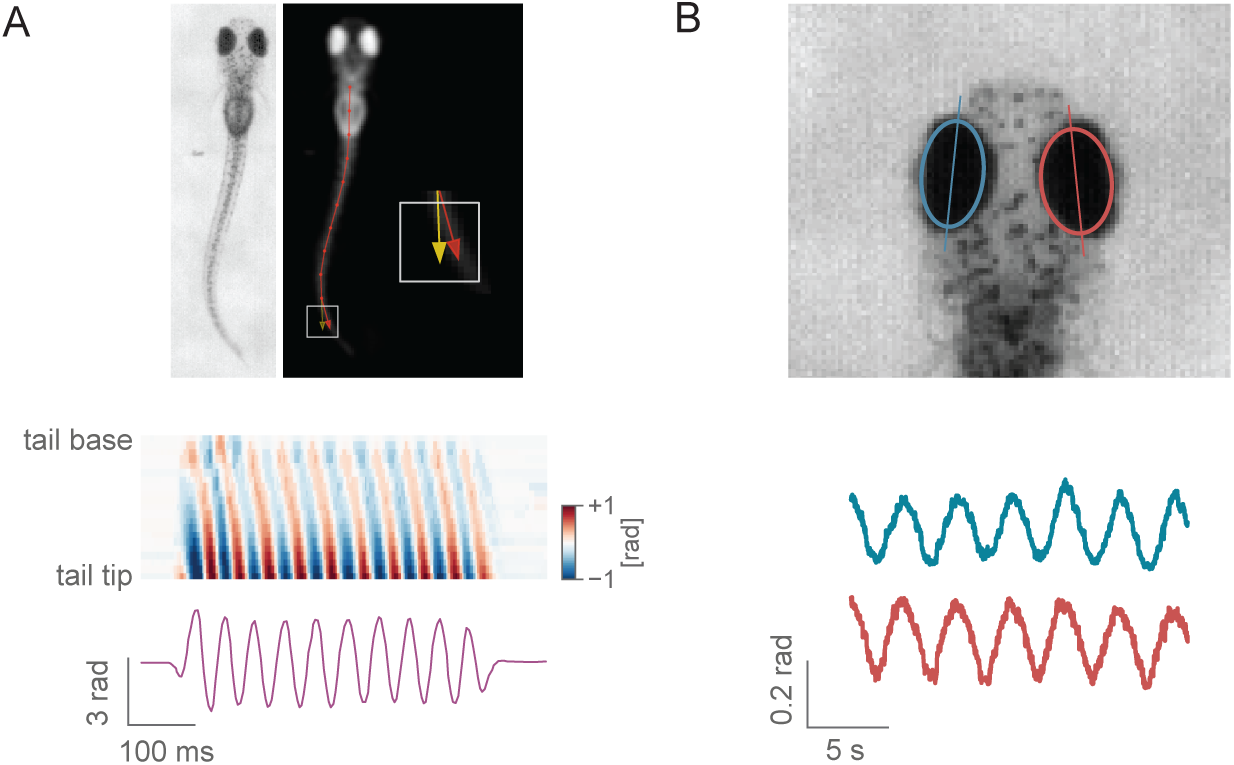
Head-restrained fish tracking in Stytra. A) Tail tracking. Above: The image is first pre-processed by inverting, downscaling, blurring and clipping, resulting in the image on the right, where the fish is the only object brighter than the background. Then, tail tracing starts from a user-defined point, and in the direction determined by another user-defined point at the end of the tail at rest. For each segment, a square in the direction of the previous segment is sampled (in red), and the direction for the next segment is chosen as the vector connecting the previous segment end and the center of mass of the sampled square (in yellow). Below: a heatmap showing the angles of the tail segments from the start to the end of the tail during a bout, and a trace representing the cumulative curvature sum from a behaving animal. The curvature sum is just the difference in angle between the first and last tail segment. B) Eye tracking. Above: eyes are detected by fitting an ellipse to the connected components of the image of the fish head after thresholding. Below: example trace of eye motion in response to a full-field rotating background.

To find the tail segments, two different functions are implemented. The first one looks at pixels along an arc to find their maximum (or minimum, if the image is inverted) where the current segment would end (as already described in e.g. [6]). On the other hand, to obtain continuous tail angles, we use a different method, based on centers of mass of sampling windows (Fig 3A). The image contrast and tail segment numbers have to be adjusted for each setup, which can be easily accomplished through the live view of the filtering and tracking results. The resulting tail shape is kept constant by interpolating a fixed number of output points regardless of the number of traced points.

##### Eye tracking

Zebrafish larvae move their eyes to stabilize the gaze in response to whole field motion, perform repositioning saccades, and converge their eyes to follow a potential pray in hunting maneuvers [7]. Naso-temporal eye movements can be described by the eye orientation with respect to the fish axis. Given the ellipsoidal shape of the eyes when seen from above, to find their orientation it is sufficient to fit an ellipse to the eye pixels and consider its axis [7]. In Stytra, a movable and scalable rectangular region can be used to select the area of the camera view containing the eyes.

As eyes are usually much darker than the background, with proper illumination conditions it is sufficient to binarize the image with an adjustable threshold which selects the pixels belonging to the eyes. Then, functions from the OpenCV library [5] are used to find the two largest connected components of the binarized region and fitting an ellipse to them. The absolute angle of the major axis of the ellipse is recorded as the eye angle (Fig 3B). A live preview of the binarized image and the extracted ellipses helps the user to adjust the parameters.

#### Freely-swimming fish tracking

To support different kinds of paradigms where fish are not embedded, we provide functions for freely-swimming fish tracking. The behavioral paradigms applicable range from investigating taxis under different environmental conditions to characterizing motion kinematics. To track the fish in an open arena, the first required step is background subtraction. The background is modelled with a mean image taken from multiple frames averaged in time, and slowly updated with an adjustable time constant. The subsequently processed image is the negative difference between the current frame and the threshold (pixels that are darker than the background are active). This image is first thresholded and regions within the right area range are found. Both eyes and the swim bladder are found as darker parts inside of these regions, and the center of mass of the three objects (two eyes and swim bladder) is taken as the center of the fish head. The direction of the tail is found by searching for the point with the largest difference from the background on a circle of half-tail radius. This direction is subsequently refined in the course of tail tracking, as described in the tail tracking section. The kinematic parameters are smoothed with Kalman filtering. An example fish tracking result is shown in Fig 4. Up to 5 fish, and with further optimizations potentially more, can be tracked at the same time. Their identities are maintained constant while they are in the field of view and not overlapping, by keeping track of the previous positions and orientations. The number of fish does not significantly impact performance, however the resolution of the camera does, so we recommend a well-configured modern computer for tracking multiple fish over larger fields of view.

**Fig 4.**
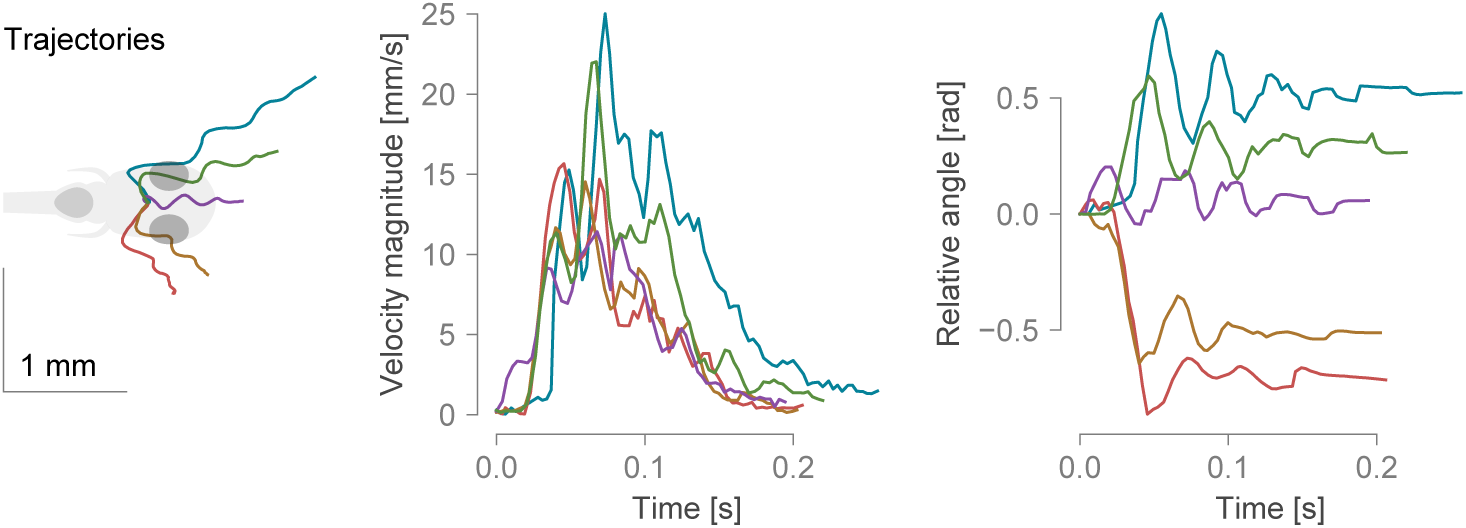
Example bouts tracked from freely-swimming fish. From left to right: trajectories of bouts in different directions, the velocity magnitude and the total angle change during the course of the bouts. The data was sampled at 300 Hz.

For closed-loop experiments, the camera view and the projected area need to be aligned. To this end, a calibration module inside of Stytra finds the mapping between the area covered by the camera and the area illuminated by the screen. During calibration, three points are projected on the screen and detected as local maxima on the camera image. Then, a transformation matrix is computed to align the projected and recorded points. If the setup elements are kept firmly in place, the calibration has to be done only once, however regular checking of the calibration on a weekly basis is encouraged.

### Closed-loop stimuli design

Stimuli whose state depends on the behavior of the fish (position and orientation for freely swimming fish, and tail or eye motion for embedded fish) are controlled by linking the behavioral state logs to stimulus display via `Estimator` objects (see Fig 1). An `Estimator` receives a data stream from a tracking function (such as tail angles), and uses it together with calibration parameters to estimate some quantity online. For example, a good proxy for fish velocity is the standard deviation of the tail curvature over a window of 50 ms [8], which we refer to as vigor. Fig 5 shows an example of how vigor can be used in a closed-loop optomotor assay. When presented with a global motion of the visual field in the caudal-rostral direction, the fish tend to swim in the direction of perceived motion to minimize the visual flow, a reflex known as the optomotor response [3, 9]. The visual feedback during the swimming bout is a crucial cue that the larvae use to control their movements. In this closed-loop experiment, we use the vigor-based estimation of fish forward velocity, together with a gain factor, to dynamically adjust the velocity of the gratings to match the visual flow expected by a forward swimming fish. The gain parameter can be changed to experimentally manipulate the speed of the visual feedback received by the larvae [8] (see below).

**Fig 5.**
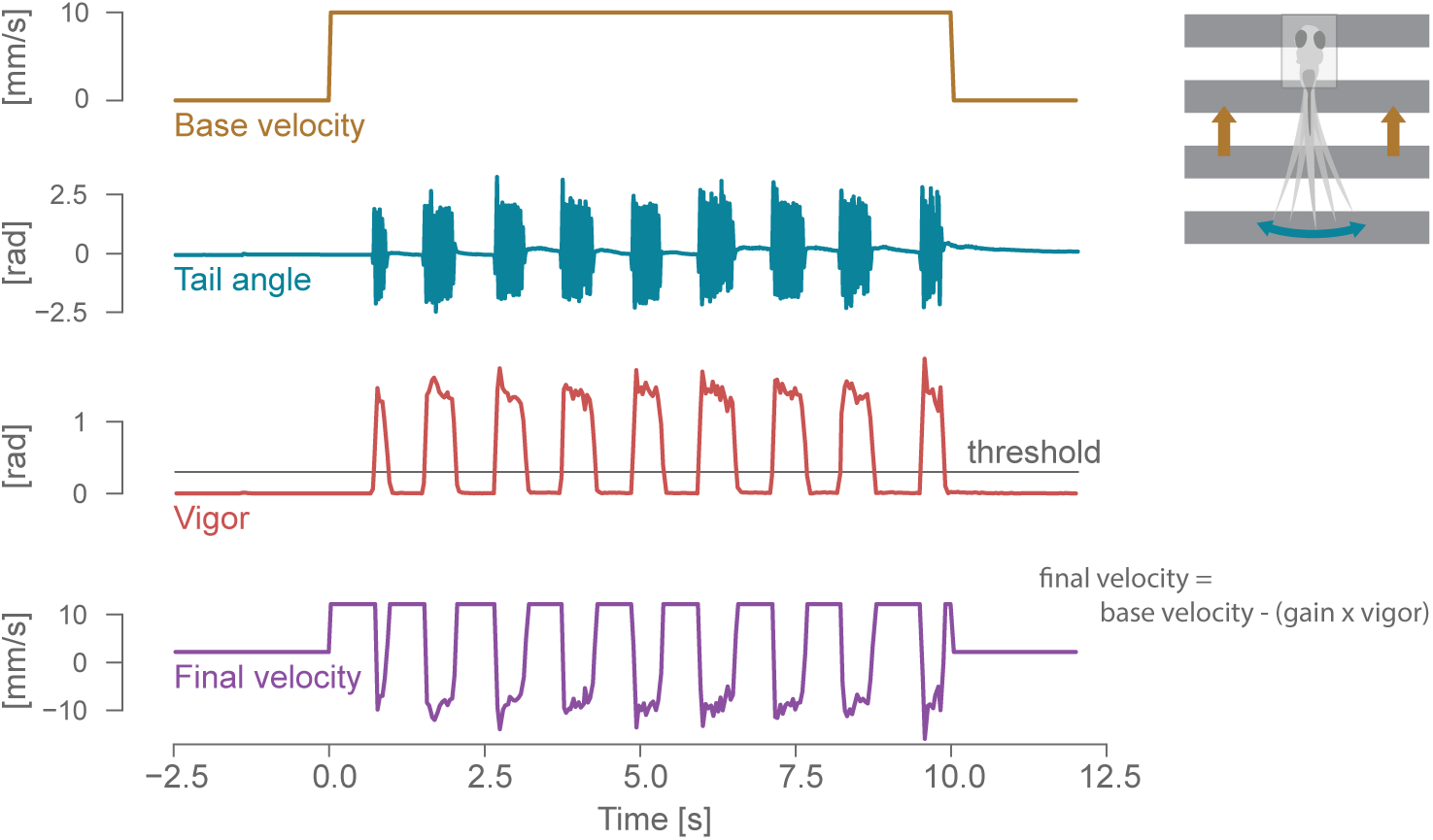
Closed-loop optomotor assay. Dynamic update of the stimulus in a closed loop assay for the optomotor response. From top: open-loop velocity of the gratings moving caudo-rostrally below the fish; cumulative tail angle (see the tail tracking section and Fig 3 for details); bout vigor, estimated by calculating the instantaneous standard deviation of the angle sum in a 50 ms window; final closed-loop velocity of the gratings, with backward movements induced by the fish tail bouts.

Closed-loop stimuli may be important for freely swimming fish as well, for example to display patterns or motion which always maintain the same spatial relationship to the swimming fish by matching the stimulus location and orientation to that of the fish.

### Synchronization with external devices

Stytra is designed to support the presentation of stimuli that need to be synchronized with a separate acquisition program, *e.g.* for calcium imaging or electrophysiology. To this end, the `Trigger` object enables communication with external devices and different computers to synchronize the beginning of the experiment. When added to an experiment, a `Trigger` ensures that the stimulation protocol does not start until a signal is received from the external device. Two ways of receiving the triggering signal are already supported in the library: TTL pulse triggering via a LabJack board, and communication over a local network employing the ZeroMQ library. Messages exchanged through ZeroMQ can also contain data such as the microscope configuration, that will be saved together with the rest of the experiment metadata. The triggering module is designed to be easily expandable, and we provide instructions for writing custom trigger objects. In our lab the two-photon microscope is controlled by custom LabView software, we expanded it to include ZeroMQ communication with Stytra. An example LabView program that can be used to trigger Stytra is illustrated in the triggering section of the documentation. In Results, we describe an example experiment using this triggering configuration to link behavioral and stimulus quantities and the recorded calcium responses. Proprietary scanning programs where this cannot be achieved can still trigger Stytra using TTL pulses.

### Data collection

The design of Stytra encourages automatic data management: all parametrized aspects of the experiment are saved by a `DataCollector` object. Various logs accompanying the experiment run (state of the stimuli, the raw tracking variables and the estimated state of the fish) are saved as tabular data. The supported data formats are CSV, HDF5 and Feather but others could be added as long as they are supported by a Python library. To demonstrate the convenience of the data and metadata saving methods of Stytra, we made example data available together with Jupyter notebooks for the analyses that can reproduce the figures in the paper. A central experiment database can be connected to keep track of all the experiments in a lab or institute. The documentation provides an example of a MongoDB database connection.

### Setup hardware

In our effort to make experiments as open and reproducible as possible, we documented example setups that can be used together with the Stytra software for performing behavioral experiments in head-restrained and freely swimming fish (Fig 6). In general, the minimal setup for tracking the fish larvae requires a high-speed camera (a minimum of 100 Hz is required, but we recommend at least 300 Hz to describe the details of the tail kinematics). The camera must be equipped with a suitable objective: a macro lens for the head-restrained tail tracking or a normal lens for the freely swimming recordings, where a smaller magnification and a larger field of view are required. More detailed camera and lens guidelines can be found in the documentation. Infrared illumination is then used to provide contrast without interfering with the animal’s perception. Since fish strongly rely on vision and many of their reflexes can be triggered by visual stimulation, the setup is usually equipped with a projector to present the visual stimulus to the fish. Although in our setups stimuli are projected below the fish, a lateral projector would be fully compatible with Stytra.

**Fig 6.**
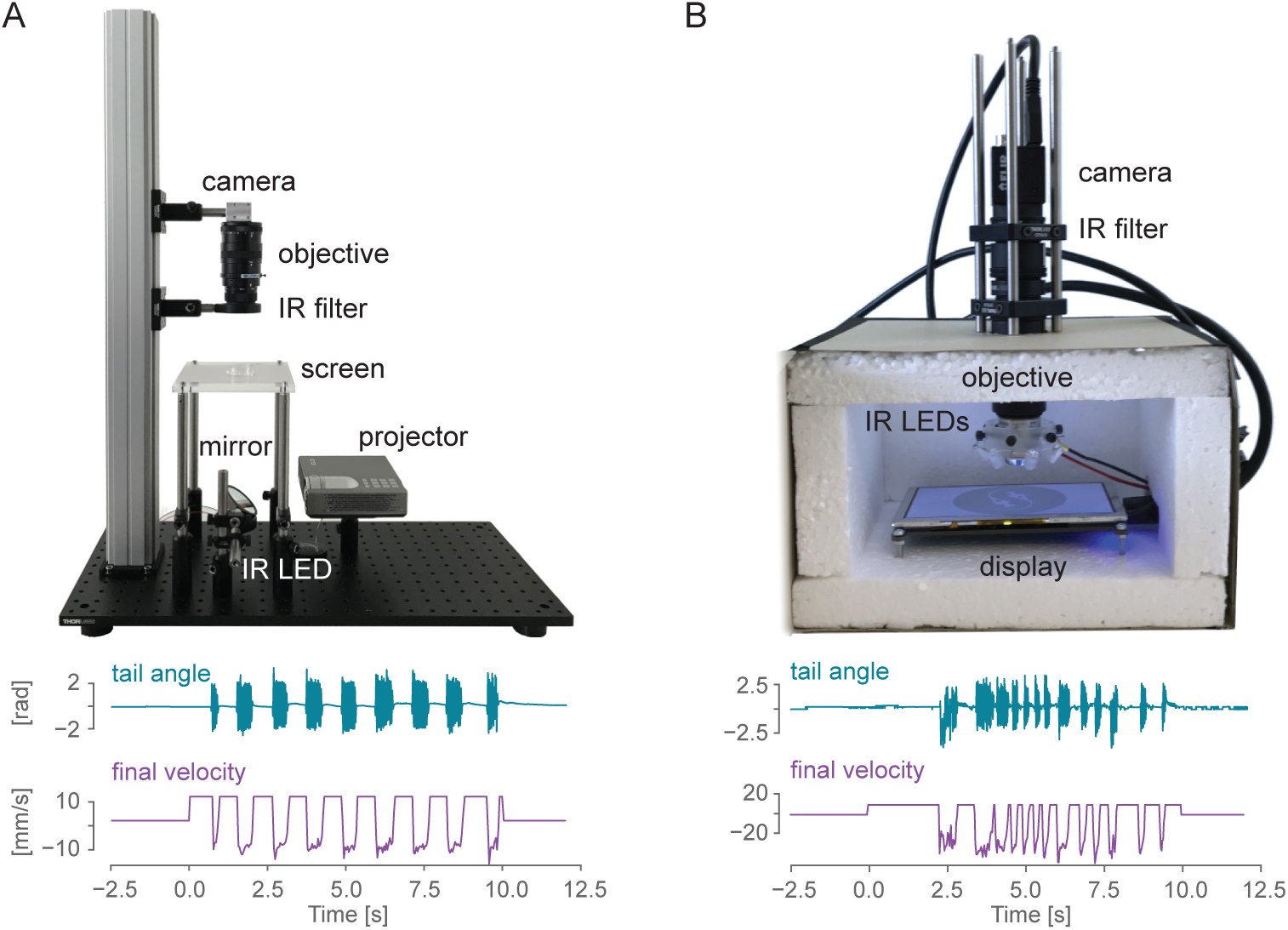
Hardware for zebrafish behavior experiments. A) Sample image of a behavioral setup that can be used to track embedded zebrafish tail end eyes (the opaque enclosure has been removed for visualization purposes). B) A low-cost version of the setup in A) that can be used to investigate behavior in the embedded fish. A detailed description of the setup together with a complete list of parts can be found at ***www.portugueslab.com/stytra/hardware*** list.

Most of our rig frames consist of optomechanical parts commonly used for building microscopes. These parts are convenient but not strictly necessary to build a well-functioning rig. Replacing them with simple hardware-store and laser-cut components can significantly reduce the costs. Therefore, we also provide instructions for a head-embedded setup build inside a cardboard box, where the most expensive item is the high-speed camera, putting the price of the whole setup without the computer under 700 euros. We created and described in the documentation such a setup, where we were able to elicit and record reliable optomotor responses in larval zebrafish (Fig 6).

A complete description of all the above-mentioned versions of the setup along with an itemized list of parts is included within the Stytra hardware documentation.

### Comparison with existing software packages

Many general-purpose systems have been proposed over the years to create (visual) stimuli and control behavioral experiments, each with its own strengths and limitations. Below we sum up some of the libraries which are currently maintained, and we present how they compare to Stytra.

#### Bonsai

One of the most widely-used frameworks is Bonsai [1], a very flexible system built in the language C# that enables defining custom acquisition and stimulation pipelines using a dataflow-based visual programming interface. In principle, all functions of Stytra can be implemented in Bonsai. However, they would require a large amount of graphical programming effort, similar to LabView. Moreover, Bonsai is written using Microsoft .NET technologies which have a limited support on non-Windows platforms. Although Python scripting is partially supported, the primary extension language is C# which far fewer scientists are acquainted with.

#### Psychophysics Toolbox

Psychophysics Toolbox [2] offers a large toolbox to build visual stimuli and stimulation protocols. The toolbox has been developed with human psychophysics in mind, in particular visual and auditory psychophysics. It provides large control over display and sound hardware, and many tools for acquiring responses from the subject through mouse and keyboard. Still, its application is restricted to the stimulus design, as it does not offer any camera integration or animal tracking modules. This makes it ill suited for developing closed-loop stimuli where behavior and responses of the animal need to be fed back to the stimulus control software. Moreover, it is written in the proprietary software package Matlab.

#### Psychopy

Psychopy [10] is a library similar to the Psychophysics Toolbox, written in Python. It provides precise control over displaying visual and auditory stimuli (not currently supported in Stytra), and a set of tools for recording responses through standard computer inputs (mouse and keyboard). However, it is also purely a stimulation library without video or other data acquisition support.

#### MonkeyWorks

MonkeyWorks is a C/C++ library to control neurophysiological experiments, developed mostly for (visual) neurophysiology in primates and rodents. It provides support for building complex tasks involving trials with different possible outcomes, and contains a dedicated library for handling visual stimuli. However, it is not designed for online analysis of video recordings of behavior, which are essential for closed-loop experiments. Moreover, while scripting and expanding Stytra requires pure Python syntax, experiments in MonkeyWorks are coded in custom high-level scripting language based on C++. Most importantly, it runs only on Mac OS, which depends on Apple hardware, used only in a minority of laboratories.

#### ZebEyeTrack

The software solution described in [11] covers a small subset of Stytra functionality - eye tracking and eye-motion related stimulus presentation. It is implemented in LabView and Matlab, which adds two expensive proprietary software dependencies. Running an experiment requires launching separate programs and many manual steps as described in the publication. The tracking frame rate is limited to 30 Hz in real-time while Stytra can run eye tracking at 500 Hz, and the performance is mainly limited by the camera frame rate.

As far as we are aware, none of the solutions described above offers fully automated data and metadata management, which is enforced by the design of Stytra.

## Results

### Triggering Stytra from a scanning two-photon microscope

We demonstrate the communication with a custom-built two-photon with a functional calcium imaging experiment. We performed two-photon calcium imaging in a 7 days post fertilization (dpf), head-restrained fish larva pan-neuronally expressing the calcium indicator GCaMP6f (Tg:elavl3:GCaMP6f, [12]). For a complete description of the calcium imaging protocol see [13]. These and following experiments were performed in accordance with approved protocols set by the Max Planck Society and the Regierung von Oberbayern.

We generated in Stytra a simple protocol consisting of either open- or closed-loop forward-moving gratings, similar to the optomotor assay described in the closed-loop section with gain either 0 or 1. At the beginning of the experiment, the microscope sends a ZeroMQ signal to Stytra, as described in the previous section. This triggers the beginning of the visual stimulation protocol, as well as the online tracking of the fish tail, with a 10-20 ms delay. To match behavioral quantities and stimulus features with their evoked neuronal correlates, we used the data saved by Stytra to build regressors for grating speed and tail motion (for a description of regressor-based analysis of calcium signals, see [6]). Then, we computed pixel-wise correlation coefficients of calcium activity and the two regressors. Fig 7 reports the results obtained by imaging a wide area of the fish brain, covering all regions from the rhombencephalon to the optic tectum. As expected, calcium signals in the region of the optic tectum are highly correlated with motion in the visual field, while events in more caudal regions of the reticular formation are highly correlated with swimming bouts. The Stytra script used for this experiment is available in `stytra/example/imaging exp.py`.

**Fig 7.**
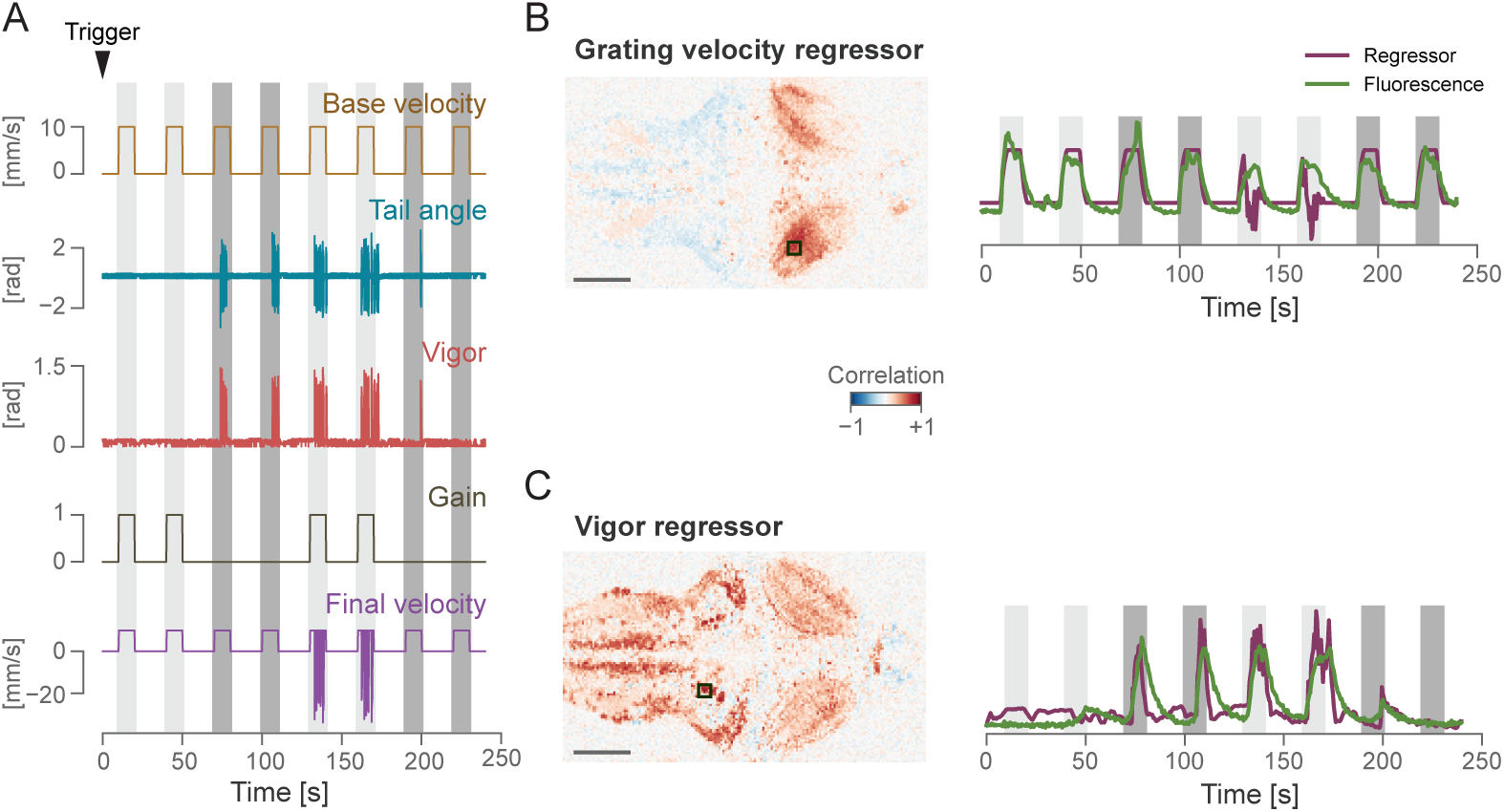
Closed loop protocol and simultaneous whole-brain calcium imaging. A) A protocol consisting of either open- or closed-loop backward-moving gratings was presented to a Huc:GCaMP6f 7 day old zebrafish larvae during two-photon imaging. The arrowhead represents the trigger signal received from the microscope. Colored stripes indicate periods when the gratings were moving: dark gray represents open loop trials (gain 0) and light gray represents closed-loop trials (gain 1). B) Left: Pixel-wise correlation coefficients with the grating velocity regressor. The square on the regressor map reports the position of the area that was used to compute the calcium trace displayed on the right. Scale bar indicates 100 um. Right: z-scored fluorescence trace from the selected area, imposed over the regressor trace. C) Same as B, for the vigor regressor.

### Experiment replication

One of the main strengths of Stytra is the possibility of sharing the experimental paradigms described in a publication as scripts that can be run on different platforms and experimental hardware. To prove the validity of this approach, we decided to showcase the software reproducing the results from two publications that investigated different behaviors of the larval zebrafish. This allowed us to verify the performance of our package in producing and monitoring reliable behavioral responses, and proved how the Stytra platform can be used to share the code underlying an experimental paradigm. The scripts used for designing these experiments are available in our repository, together with a full list of parts and description of the hardware. In this way, everyone can independently replicate the experiments simply by installing and running Stytra on a suitable behavioral setup.

#### Closed-loop motor adaptation

To demonstrate the effectiveness of the closed-loop stimulation software for head-restrained larvae, we re-implemented in Stytra one of the paradigms described in [8]. This paper addresses the importance of instantaneous visual feedback in the control of the optomotor response in 7 dpf zebrafish larvae.

In [8] closed-loop paradigm was used to have real-time control over the visual feedback that the animal receives upon swimming. After triggering motor activity with forward-moving black and white gratings (10 mm/s, 0.1 cycles/mm), online tail tracking was used to estimate the expected velocity of the fish based on freely-moving observations and a backward velocity proportional to the expected forward velocity was imposed over the forward grating speed. In one crucial experiment (Fig 3 of [8]) the authors demonstrated that reducing or increasing the magnitude of this velocity by a factor of 1.5 (high gain) or 0.5 (low gain) resulted in modifications of the bout parameters such as bout length and inter-bout interval (temporal distance between two consecutive bouts). Fig 8A shows the inter-bout interval along the protocol, where the three gain conditions were presented in a sequence that tested all possible gain transitions. When gain increased the fish was consistently swimming less (higher inter-bout interval), and *vice-versa* when gain increased. Therefore, as expected, fish adapted the swimming parameters to compensate for changes in visual feedback.

**Fig 8.**
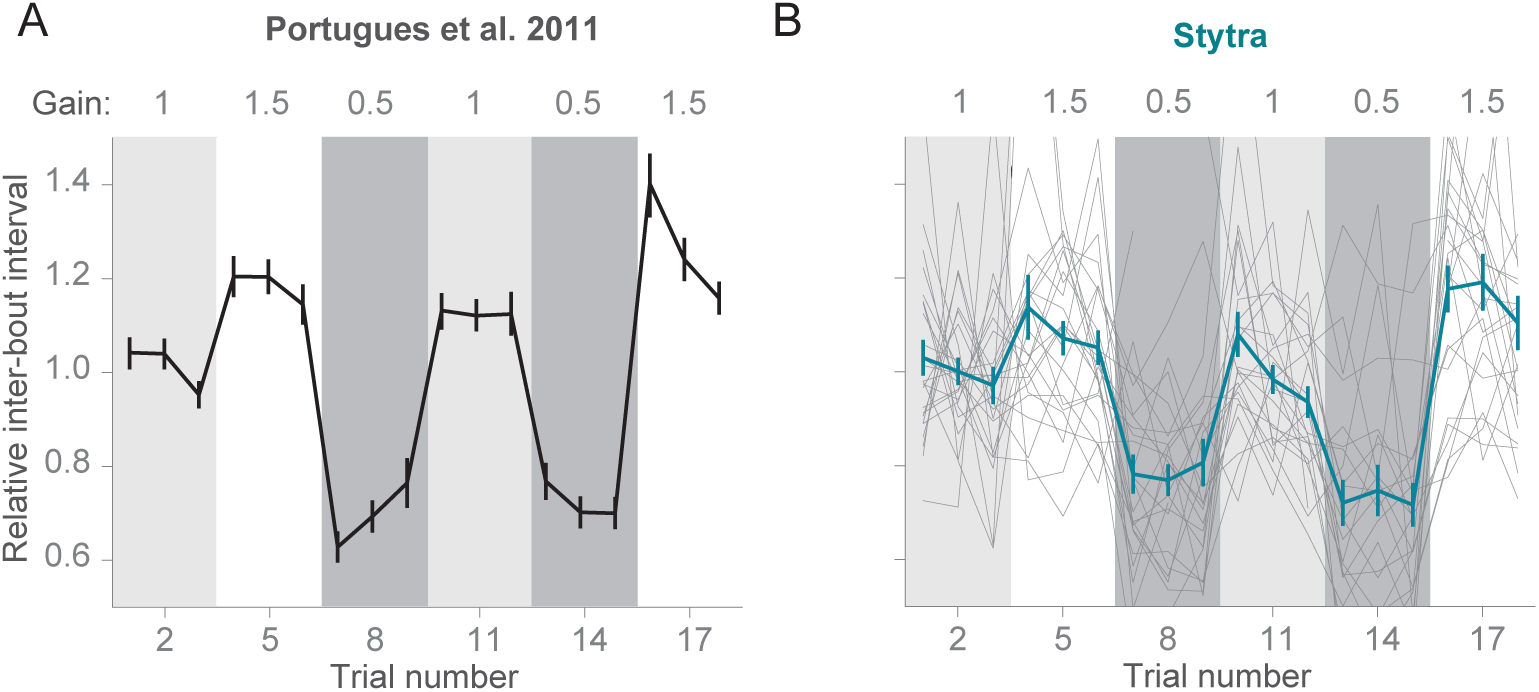
Visual feedback changes inter-bout interval in an embedded optomotor assay. Replication within Stytra of the results published in [8]. A) Changing the gain used to convert the fish vigor to backward gratings velocity affects the inter-bout interval. Line represents average interbout time normalized to the average, bars represent SEM from n=28 larvae (adapted from [8]). B) Replication in Stytra of the same experimental protocol. Line represents average interbout time normalized to the average, bars represent SEM from n=24 larvae. Individual results from each fish are represented in gray.

We reproduced exactly the same protocol within Stytra, and we used Stytra modules for closed-loop control of a visual stimulus to compare whether it could replicate the findings from [8]. Cumulative angle of the extracted tail segments were used with a gain factor to estimate fish velocity as described in [8]. The gain factor was changed in a sequence matching the protocol in [8]. The replication with Stytra yielded the same result (Fig 8B), that inter-bout interval decreased in low gain conditions and increased in high gain conditions.

#### Closed loop phototaxis assay

To test the freely swimming closed-loop performance, we replicated the protocol from [14]. The fish is induced to perform phototaxis by keeping half of its visual field bright while the other is dark. The fish is more likely to turn to the bright side. The stimulus is constantly updated so that the light-dark boundary is always along the midline of the fish. We replicated the qualitative trends observed in [14], however the ratios of forward swims to turns are notably different (Fig 9). The variability of fish responses and differences in the stimulus presentation setup (e.g. projector brightness) could contribute to these differences. Also, to reduce duration of the experiments, we included a radially-inward moving stimulus that brings the fish back into the field of view.

**Fig 9.**
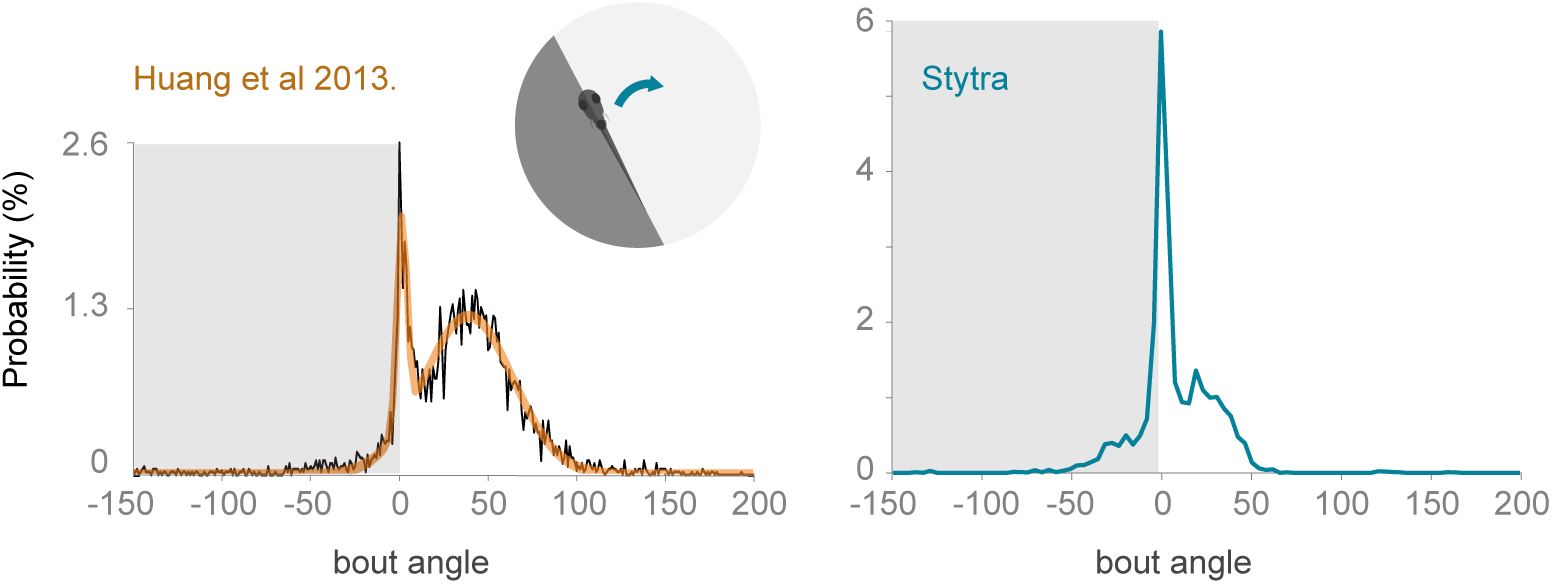
Comparison of turning angle distribution in a closed-loop freely-swimming phototaxis experiment. Left: a histogram of the angle turned per bout, redrawn from [14] Right: the equivalent panel, with n=10 fish and the protocol run with Stytra. The dark shading on the plot represents the dark side of the visual field.

## Conclusions

Our software is capable of combining online behavioral analysis with imaging of neural activity and reproducing behavioral results from established experimental paradigms. This proves its suitability as a framework for coding and running experiments in systems neuroscience. In addition to the open-source software, we are contributing to the nascent open hardware movement [4] and are providing a complete description of the hardware used for conducting behavioral experiments. Finally, we provide a set of example analysis scripts for the experiments described in this manuscript, which can be easily modified for other experimental questions.

The current version of the software supports all experimental paradigms currently running in our lab. Other setups with multiple screens or cameras would require some extensions in the architecture. Also, obtaining millisecond-level or higher temporal precision in stimulation, for optogenetics for example, would require rewriting the stimulation module in a language other than Python.

The modular and open-source nature of the package (licensed under the GNU GPL v3.0 licence) facilitates contributions from the community to support an increasing number of hardware devices and experimental conditions. Moreover, the simplicity of the implementation of an experiment within Stytra facilitates the collaboration between laboratories, since complex experimental paradigms can be run and shared with simple Python scripts. We will also make use of the community features of Github to provide support in adopting the package in other labs. Althugh Stytra has been developed with zebrafish neuroscience in mind, it can provide a good foundation for Python-based behavioral experiment control in other animals. In conclusion, we hope that Stytra can be a resource for the neuroscience community, providing a common framework to create shareable and reproducible behavioral experiments.

## Online resources

- stytra repository: https://github.com/portugueslab/stytra
- stytra documentation: http://www.portugueslab.com/stytra/
- data analysis notebooks: https://github.com/portugueslab/example stytra analysis
- example data from stytra: https://zenodo.org/record/1692080

## Author contributions

V. Š. and L.P. designed the architecture of the software and developed all functions. A.M.K. provided initial versions of camera connection and eye-tracking code, as well as ideas for the setups. R.P designed initial versions of the hardware setups and provided input for designing the tracking functions. L.P. and V. Š. performed all the experiments, and V. Š., L.P. and R.P. wrote the manuscript with input from A.M.K..

## Acknowledgments

We thank Marco Albanesi for testing the software and the first pull requests, and Virginia Palieri, Elena Ioana Dragomir and Ot Prat for being the first users of Stytra in the lab. We thank the Python open-source community on whose work this package is based on, especially Luke Campagnola for developing the invaluable PyQtGraph package. RP was funded by the Max Planck Gesellschaft (www.mpg.de) and by the Human Frontier Science Program (www.hfsp.org) via grant RGP0027/2016.

